# High-fat diet impacts the colon and its transcriptome in a sex-dependent manner that is modifiable by estrogens

**DOI:** 10.1101/2020.06.03.131771

**Authors:** L. Hases, A. Archer, R. Indukuri, M. Birgersson, C. Savva, M. Korach-André, C. Williams

## Abstract

Epidemiological studies highlight a strong association between obesity and colorectal cancer (CRC), especially in men. Estrogen, on the other hand, is associated with protection against both the metabolic syndrome and CRC. The colon is the first organ to respond to a high-fat diet (HFD), and estrogen receptor beta (ERβ) in the intestine appears to prevent CRC. How estrogen impacts the colon under HFD condition has, however, not been investigated. Estrogen can act through three different receptors (ERα, ERβ, GPER1) which all may impact metabolism. In an effort to dissect this, we fed mice a control diet or a high-fat diet (HFD) for 13 weeks and administered receptor-selective estrogenic ligands for the last three weeks. We recorded corresponding physiological impact on fat distribution, fasting glucose, colon crypt proliferation and immune cell infiltration, and the colon transcriptome response. We identify clear sex-differences at the transcriptome level, both at base line and after HFD and ligand treatments. An unexpected observation was the significant sex-differences and impact by HFD and estrogens on circadian clock gene expression, such as *Npas2* and *Arntl (Bmal1)*, in the colon. Both sexes also exhibited an increased infiltration of F4/80+ macrophages as a result of HFD. In males, but not females, this was accompanied by changes in colonic epithelial cell proliferation. ERα-selective PPT treatment had significant systemic effects, reducing body weight in both sexes, whereas ERβ-selective DPN treatment did not impact body weight, but reduced infiltration of F4/80+ macrophages in colon of both sexes and attenuated HFD-induced proliferation of male colon crypts. Both ERα and ERβ activation contributed to circadian clock gene regulations. We detail for the first time how HFD and estrogens modulate the colon transcriptome and physiology in a sex and ER-specific manner.

## Introduction

Obesity represents a worldwide public-health issue and epidemiological studies highlight a strong association between body mass index (BMI) and risk of colorectal cancer (CRC) (1-4). A high-fat diet (HFD) increases this risk, but exactly how is not clear. It is known that inflammatory bowel disease (IBD) increases the risk of CRC, and that both obesity and HFD induces a general low-grade inflammatory state (5, 6). The colon is the first organ to respond to HFD (7), with reported increase in inflammation, PPARδ signaling, stem cell activity, and altered gut microbiota (8-12), which may all contribute to initiation of CRC (7, 9, 13, 14). Interestingly, while obesity is a risk factor for both sexes, this association appears stronger in men (15). Men also have a higher incidence of colon and rectal cancers (16, 17). A role for sex hormones is supported by epidemiological studies (reviewed in Lobo *et al*.), and especially estrogen appears protective against CRC (18-20). Numerous clinical studies have also indicated a protection by estrogen against several aspects of obesity and the metabolic syndrome (21, 22), and hormones also impact the immune system (reviewed in Boman *et al*. (23)).

The biological action of estrogen is mediated by three receptors, the nuclear receptors ERα (ESR1) and ERβ (ESR2), and the transmembrane G protein-coupled estrogen receptor 1 (GPER1). These receptors are expressed in both women and men, with a tissue-specific expression pattern. Both ERα and ERβ have been implicated to elicit anti-obesogenic effects (24-26). While both receptors may protect against HFD-induced obesity, ERα clearly improves the metabolic profile, whereas the role of ERβ in this respect is controversial. Most of the CRC protective effect, however, has been linked to ERβ. Treatment with ERβ-selective agonist, for example, resulted in anti-inflammatory and anti-tumorigenic effects in different CRC mouse models (27-30), and loss of ERβ results in pro-inflammatory environment and increased tumor development in both APC ^Min/+^ and colitis-associated tumor mouse model (31, 32)(Hases et al, under review). Notably, intestinal-specific deletion of ERβ strongly enhanced inflammatory signaling in the colon of both male and female mice (Hases et al, under review). However, the impact of ERβ in modulating the colon inflammation during HFD-induced obesity has not been explored. We hypothesized that estrogenic signaling, trough ERα, ERβ or both, can attenuate the effect of HFD in colon, with possible sex-differences.

In order to test this, and to characterize the impact of estrogen signaling on the colon microenvironment under HFD, we performed a detailed analysis of mice of both sexes which were fed either a control diet (CD) or a HFD for 13 weeks, and administered different estrogenic ligands for the last 3 weeks prior to sacrifice. Importantly, by analyzing corresponding effects on the full colon transcriptome, we generated an unbiased overview of alterations mediated by HFD and estrogens. We observed sex-differences in the colon transcriptome and in its response to HFD. We demonstrate for the first time that estrogen signaling is indeed protective against HFD-induced colonic alterations. DPN-activated ERβ restored the impaired expression of the core clock gene *Npas2* in the colon and decreased the HFD-induced epithelial cell proliferation in males. Besides, HFD-feeding induced colonic F4/80+ macrophage infiltration in both sexes, and activation of ERβ decreased macrophage infiltration. Altogether, our data show for the first time that estrogen signaling can attenuate HFD-induced alterations of the colon microenvironment, and we show that some of this effect is mediated by ERβ expression in the colon. These findings open up new insights into prevention of obesity-associated CRC.

## Results

### Base-line colon transcriptome and sex differences

Female mice had significantly higher circulating estradiol levels compared to males when fed CD, however male and female mice fed CD showed no significant sex differences in body weight (BW) gain during the study period, nor in fasting glucose levels at the study end point (Fig. 1A and B). Females showed a trend of higher percentage of total fat (TF) in relation to BW, but this was not significant (Suppl. fig. S1A). However, as also noted previously (33), the fat distribution was sex dependent. Males had more visceral white adipose tissue (VAT) and less subcutaneous white adipose tissue (SAT) in relation to total fat (Suppl. fig. S1A). As a consequence, males had a substantially lower SAT/VAT ratio (0.32 versus 0.52 in females, Fig. 1B). A low SAT/VAT ratio has been correlated to poor metabolic outcomes. To specifically study sex differences in the colon, we performed RNA-Seq of distal colon tissue from males (n=5, from two different cages) and females (n=6, from three different cages). As evident from the principal component (PCA) plot, the colon transcriptomes exhibited clear sex differences (Fig. 1C, blue circle). Gene expression analysis identified 1564 significantly differentially expressed genes (DEG) between male and female colon. In order to explore which types of genes were responsible for these sex differences, we performed enrichment analysis for biological processes (BP). The BP enrichment analysis revealed sex differences in gene expression related to immune response (e.g. *Cd36, Ccl6* and *Ccl20*), cell proliferation (e.g. *Hif1a, Igf1* and *Lgr5*) and canonical Wnt signaling (e.g. *Ccnd1, Gli1* and *Wnt2b, Fig*. 1E and F). Although the mice had been sacrificed at the same time of the day, we found sex differences in expression of circadian rhythm genes, including the clock genes *Arntl, Npas2, Cry1 Per3* and *Nr1d1/Rev-ErbA* (Fig. 1E and F). Thus, in mice fed a normal control diet, that lacks phytoestrogens, we report, for the first time, clear sex differences in the colon transcriptome.

**Figure 1.**
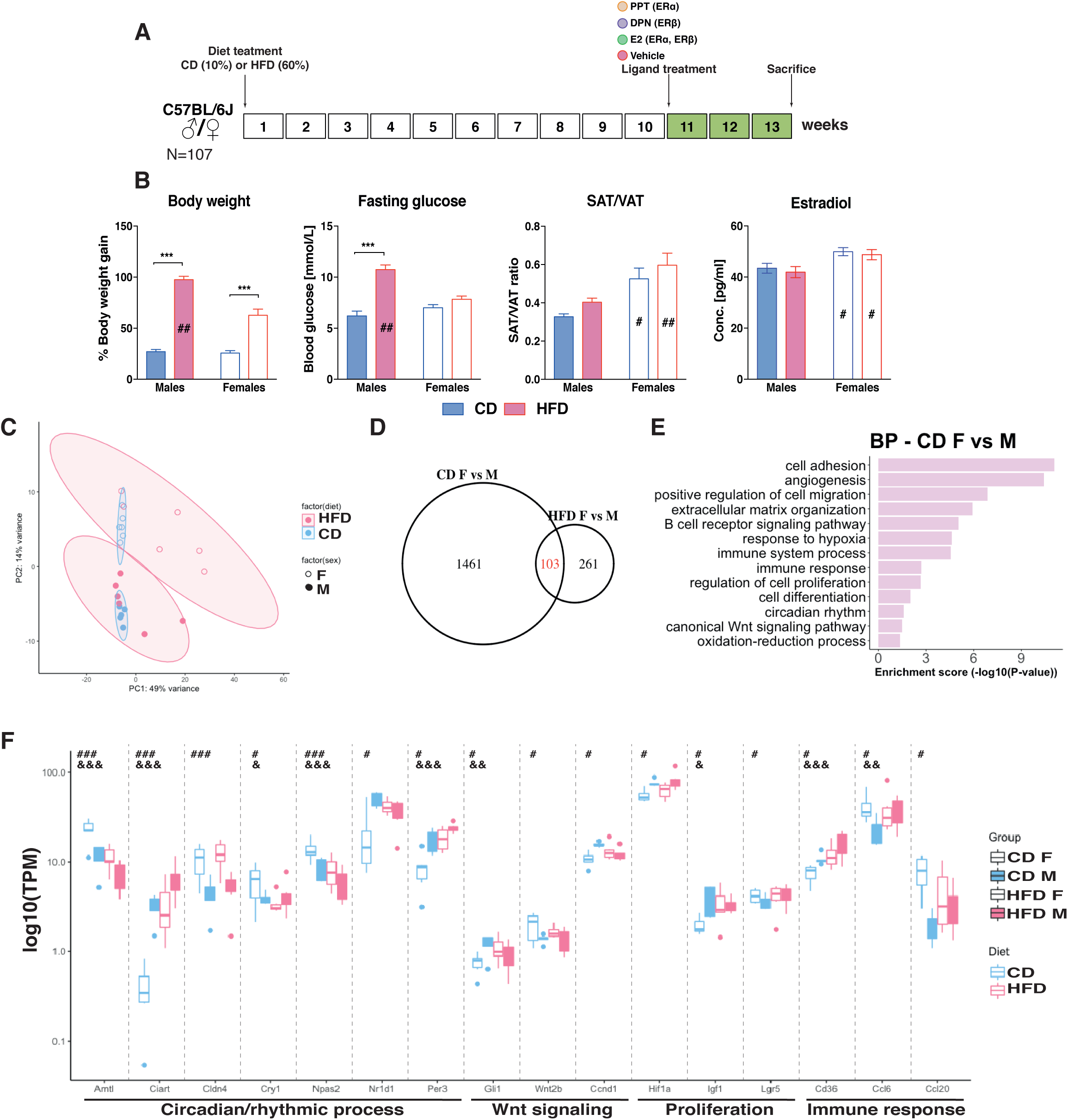
The male metabolic profile is more sensitive to HFD and there is a sex-specific gene expression profile independent of diet. (A) Experimental study design, mice of both sexes were given a CD or a HFD for 13 weeks and injected with different estrogenic ligands for the last 3 weeks prior to sacrifice. (B) Percentage of BW gain, fasting glucose, SAT/VAT ratio and serum estradiol levels. (C) Principal component analysis of RNA-seq samples for male (unfilled) and female (filled) mice fed a CD (blue) or HFD (red) (n=5-6). (D) Venn diagram comparing differentially expressed genes in colon between male and female mice fed a CD or a HFD. Genes were considered as differentially expressed with p-value<0.05 and log_2_FC>|0.4|. (E) Biological process enrichment analysis based on genes differentially expressed between the sexes during CD. (F) Boxplot of TPM values of genes differentially expressed between sexes and/or dietary conditions. The results are presented as mean ±SEM. Two-way ANOVA followed by fisher’s LSD test, * p<0.05, ** p<0.01, *** p<0.001. # Indicate significantly sex-differences during CD, ## during HFD and ### during both dietary conditions. & Indicate significantly difference between CD and HFD in females, && in males and &&& in both sexes.

### Sex differences in high-fat diet-fed mice

In order to study how the colon was affected by HFD, male and female mice were compared following 13 weeks of HFD. Female mice presented significantly higher circulating estradiol levels during HFD and HFD induced weight gain in both sexes, but significantly more in males (100% gain) than in females (65% gain, Fig. 1B). The increases of total fat mass as fraction of BW was similar in both sexes and the noted sex differences in VAT and SAT under CD remained relatively unchanged under HFD (Fig. 1B and Supp. fig. S1A). We observed that the obese males presented a significantly increased blood glucose level (after 6-h fasting), which was not observed in females (Fig. 1B). These results support previous observations that obese males are more susceptible to glucose intolerance and insulin resistance than females (33, 34). Next, in order to detail the impact of HFD on the colon, we compared the RNA-Seq data of distal colon samples for both male and female mice fed CD with those fed HFD. The PCA plot indicates that while the CD colon transcriptome clustered tightly together, the HFD one was more spread out (Fig. 1C). As a result, the sex difference under HFD was less distinct, but analysis of differentially expressed genes still revealed 364 DEGs. Over 1/4 (28%) of the sex differences under HFD was detected also under CD conditions (Fig. 1D). The most overrepresented genes among these stable DEGs (103) belonged to the circadian/rhythmic pathway (e.g. *Arntl, Ciart, Cldn4* and *Npas2*, Fig. 1F).

### HFD modulates the colon transcriptome differently in male and female

In order to explore how HFD impacted male and female colon, we compared the transcriptomes under HFD with corresponding expression under CD, for male and females separately. We identified differential expression of 568 genes in females and 632 in males (Fig. 2A). We noted that a larger proportion was upregulated in females (85%) than in males (51%), and that females exhibited a stronger upregulation. Only 75 genes responded to HFD in both males and females (Fig. 2B). Among these common genes were, again, those related to the circadian rhythm (*Arntl, Ciart, Npas2* and *Per3*, Fig. 1F), oxidation-reduction process, and inflammatory response (e.g. *Arg1* and *Anxa1*, Fig. 2A). While most of the oxidation reduction-related genes were upregulated in both females and males, *Alox15* was downregulated by HFD in males but upregulated in females (Fig. 2A). We analyzed the sex-specific transcriptomic responses to HFD for BP enrichment. Both sexes altered expression of genes with functions in cell adhesion, cell proliferation, angiogenesis, migration and immune system (Fig. 2C), but different genes were altered in males and females. Pathways involved in cell cycle (e.g *Stk11* and *Wee1*), hypoxia (e.g. *Actn4* and *Sod3*) and glucose homeostasis (e.g. *Stk11*) were particularly enriched in males, whereas regulation of apoptosis (e.g. *Nr4a1*), inflammatory response (e.g. *Cxcl5*), MAPK cascade (e.g. *Igf1*) and Wnt signaling (e.g. *Sox9*) was enriched in females (Fig. 2A and C). All together our data demonstrate for the first time that sex impacts the gene expression profile and its response to HFD in colon. While both sexes presented alterations within similar biological functions, including the immune system and cell adhesion, other changes appeared tied to sex, such as a more pronounced changes of cell cycle and hypoxia pathways in males, and of the inflammatory response and Wnt signaling in females.

**Figure 2.**
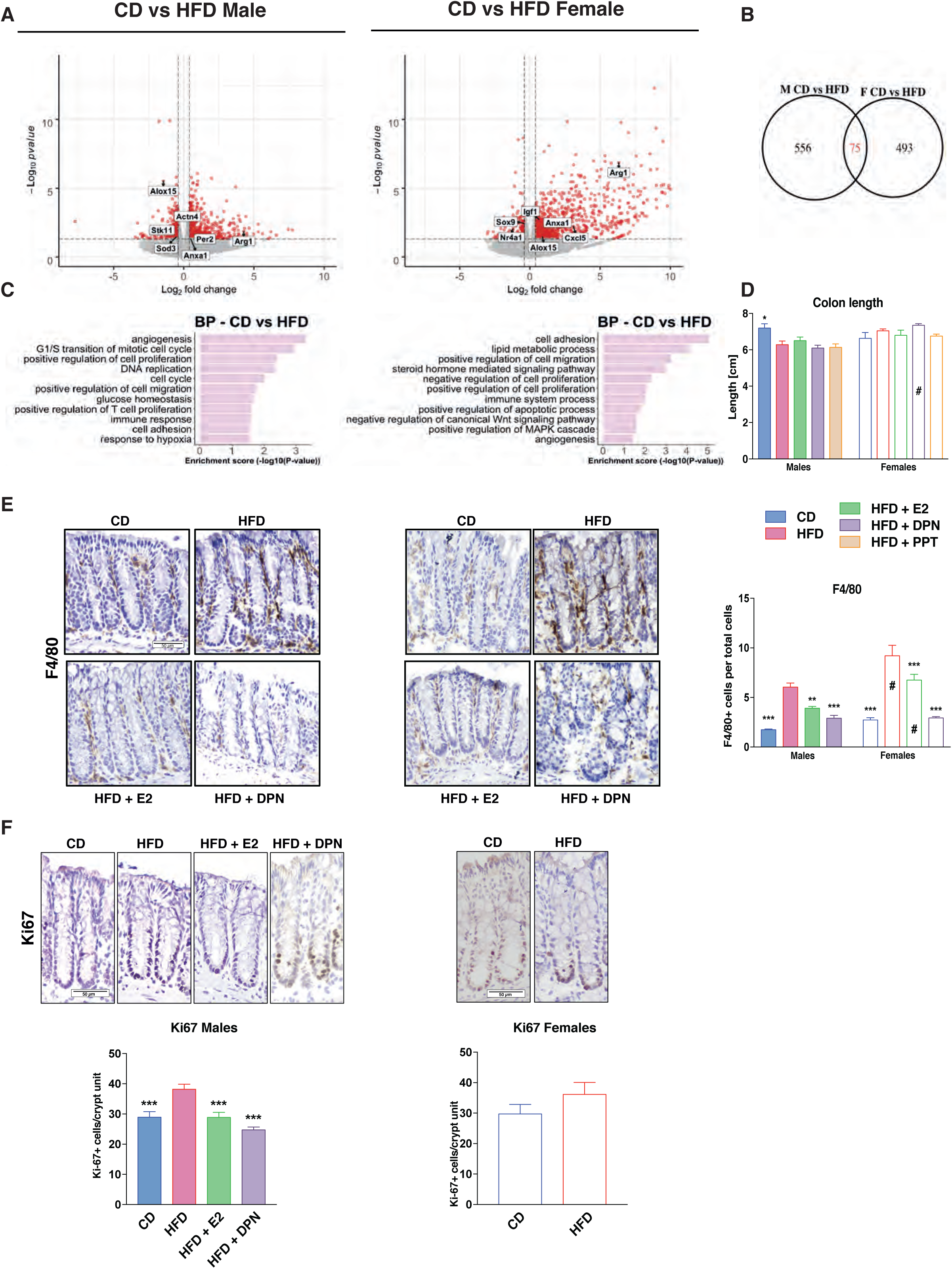
The colon transcriptome responds to HFD in a sex-dependent manner. (A) Volcano plot showing HFD regulated genes in males to the left and females to the right. Genes were considered as differentially expressed with p-value<0.05 and log_2_FC>|0.4|. (B) Venn diagram comparing differentially expressed genes in colon between CD and HFD in male and female mice. (C) Biological process enrichment analysis based on the sex-specific regulated genes in males and females. (D) Colon length measured in cm. (E) Macrophage infiltration measured with F4/80 IHC with quantification of the ratio of F4/80^+^ per total cells. (F) Intestinal epithelial cell proliferation measured with Ki67 IHC and with quantification of Ki67^+^ cells per crypt unit in males and females on a CD, HFD and HFD treated with E2 (n=5-15). The results are presented as mean ±SEM. One-way and two-way ANOVA with uncorrected Fisher’s LSD test, *p<0.05, **p<0.01, ***p<0.001. # Indicate sex-differences.

### HFD increases colon epithelial inflammation and proliferation in males

The above colon transcriptome analysis indicated that the immune system was regulated by HFD and that, especially in males, the cell cycle and hypoxia was modulated. A shortened colon is an indication of colonic inflammation, and we found that males fed a HFD presented a significantly shorter colon compared to those fed CD (Fig. 2D). This was not noted for females (Fig. 2D). Next, we explored markers for macrophages in the histological sections using immunohistochemistry. We found that mice of both sexes exhibited a significantly increased level of infiltrating macrophages when fed HFD, as measured by F4/80 staining (Fig. 2E). In addition, the males but not females presented an increased proliferation of crypt cells after HFD, as measured by Ki67 staining (Fig. 2E). This reveals that HFD results in clear sex-dependent functional effects in the colon, in accordance with the above noted transcriptional differences.

### Estrogen improves the metabolic profile of obese males via ERα

Several studies have shown that estradiol (E2) treatment improves the metabolic profile of HFD-fed rodents of both sexes (35, 36). Further, knockout of ERα results in obese mice (24). In our study, mice treated with E2, ERα-selective ligand PPT, or vehicle for the last 3 weeks (Fig. 1A) did not exhibit any differences in BW-gain in females (Fig. 3A & Suppl. fig. S1C-D). However, male mice showed a dose-dependent BW loss during treatment, with a 5% weight reduction on higher-dose E2 treatment (0.5 mg/kg BW, Fig. 3A & S1B). This was accompanied by a significant reduction of total fat (as proportion of BW) and decreased blood glucose level upon fasting, not seen in females (Fig. 3A & Suppl. fig. S1C). The higher dose E2 did not yield in unphysiological estradiol concentrations in the serum (Fig. S1B). These effects were also evident with the selective ERα ligand PPT (Fig. 3A). Thus, our experimental set-up confirms that exogenous E2 via ERα impacts obesity, fasting glucose, total fat and VAT ratios in males.

**Figure 3.**
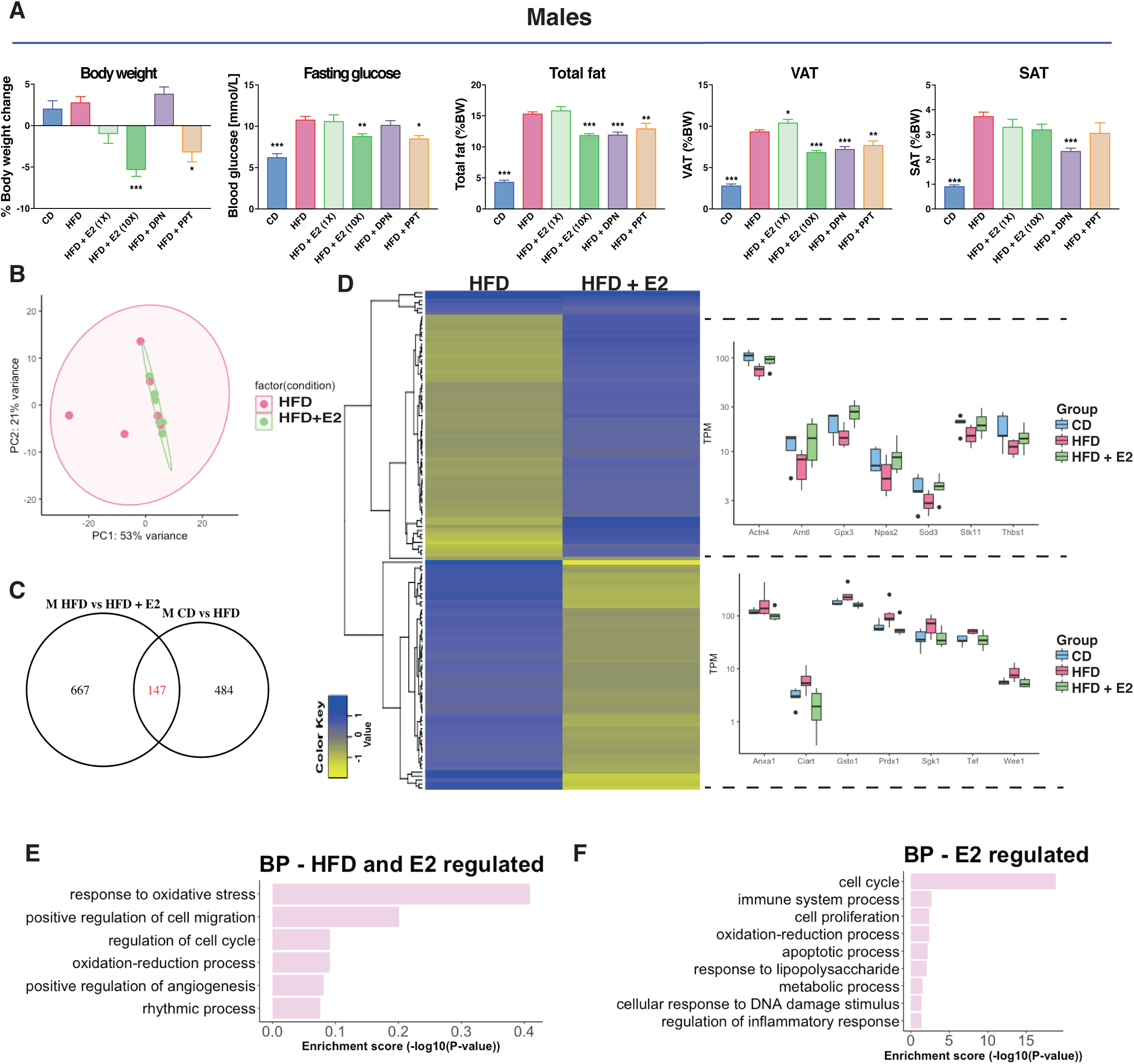
Estrogen treatment improves the metabolic profile in males. (A) BW change during treatment, fasting glucose levels, total fat mass (reported to BW) and VAT and SAT (reported to BW) for males on a CD, HFD and HFD treated with E2, DPN and PPT. (B) Principal component analysis of RNA-seq samples for males fed a HFD with and without E2 treatment. (C) Venn diagraming comparing HFD regulated genes in males with E2 regulated genes in males under a HFD. (D) Heatmap of the homogenously regulated common genes (regulated by both HFD and E2 under a HFD) and boxplots of TPM values of some genes. Biological process enrichment analysis of the (E) HFD-induced genes opposed by estrogen signaling and (F) estrogen regulated genes under HFD feeding.

### Estrogen modulates expression of specific gene sets in HFD colon

Next, we wanted to determine if E2 treatment impacted the colon, and specifically if it could modulate the HFD-induced colon inflammation signature. We focused on males, who had exhibited a clear shortening of the colon upon HFD, and had shown a significant E2-mediated impact on the metabolic profile. The colon RNA-Seq data, visualized in a PCA plot, show that the group of HFD-fed male mice treated with E2 clustered tighter together compared to vehicle treated (Fig. 3B), just as CD diet samples did (Fig. 1C). Analyzing differential gene expression, we found that E2 treatment modified the expression of 814 genes in distal colon. Of these, 147 genes overlapped with those impacted by HFD (compared to CD), and nearly all (97%) were regulated in the opposite direction (Fig. 3C-D). Two homogenous (regulated similarly in at least three animals from each group) clusters of DEG appeared, illustrated by a heatmap in Figure 3D. One cluster (58 genes) was upregulated by HFD (compared to CD), and downregulated by E2 (under HFD conditions), and comprised genes such as *Anxa1, Ciart, Gsto1, Prdx1, Sgk1, Tef* and *Wee1* (Fig. 3D, lower right). The second cluster included 61 genes that were downregulated by HFD and then upregulated by E2, such as *Actn4, Arntl, Gpx3, Npas2, Sod3, Stk11* and *Thbs1* (Fig. 3D, upper right). BP enrichment analysis revealed that genes regulated by both HFD and E2 were involved in cell cycle and rhythmic process (Fig. 3E). Most (667 of 814) E2-regulated genes were, however, not modulated by HFD. Among these, we note enrichment of cell cycle and cell proliferation (e.g. *Cdk1, Melk, Mki67*), and immune system process (e.g. *Alcam, Irf7* and *Bcl6*) functions (Fig. 3F). This demonstrates that both sex and estrogen have a large impact on the colon transcriptome, and that estrogen clearly opposes a fraction of the HFD-induced expression.

### Estrogen impacts immune cell recruitment and proliferation in the colon

Since we noted that E2 treatment could attenuate signs of inflammation and proliferation at the gene expression level, we investigated whether E2 impacted the length of colon or infiltration of macrophages. The length of the HFD colon was not significantly elongated by E2 (Fig. 2D). Macrophage infiltration was, however, clearly reduced upon E2 treatment in both male and females (Fig. 2E). We also noted a significant reduction of positive Ki67 stained cells after E2 treatment in males (Fig. 2F). These data clearly demonstrate that E2 impacts the colon, both in terms of immune cell recruitment and colonic epithelial cell proliferation.

### ERβ-selective agonist modulates fat distribution, immune cells, and cell proliferation

While ERα impacts obesity and the metabolic syndrome, it is unclear whether the effects in colon is a secondary effect of this, or if there are local effects mediated by estrogen within the colon. We have previously shown that intestinal epithelial ERβ impacts inflammation and colon carcinogenesis. To investigate effects by ERβ, we had included treatment with ligand selective for ERβ (DPN) in our animal experiment (Fig. 1A). DPN treatment did not result in weight reduction, in accordance with data assigning this phenotype to ERα. On the contrary, DPN-treated males and females exhibited a non-significant trend of increased weight gain, which was accompanied by a significant increase of blood glucose levels in females but not males (Fig. 3A & Suppl. fig. S1C). This also demonstrates receptor selectivity of the ligands in the concentrations used in this set up. Although the effects by DPN on weight were nonsignificant in males, we noted clear reductions of total fat and VAT (as percentage of BW) by DPN treatment, in a scale similar or higher than what was achieved with PPT treatment. Further, a reduction in SAT was specific for ERβ ligand activation (Fig. 3B). We did not see these effects in females. This suggests that while obesity and HFD-induced BW can be decreased by ERα, ERβ appear to modify the ratio of total fat, VAT, and SAT in males. In addition, we saw that estrogen through ERβ could oppose the HFD-induced Ki67 proliferation in males (Fig. 2F). The increased F4/80 macrophage infiltration upon HFD was significantly reduced by estrogen through ERβ in both sexes (Fig. 2E). Our data establish that while the improved metabolic profile is mediated via activation of ERα, ERβ activation impacts the colon by attenuating HFD-induced colonic proliferation in males and macrophage infiltration in both sexes.

### ERβ regulates gene expression in the colon

As shown above, estrogen treatment (0.5 mg/kg BW) opposed a set of HFD-induced colonic gene expression in males. This could be an indirect consequence of systemic effects (e.g. ERα improving the overall metabolism) or direct local effects through estrogen receptors expressed in the colon. We have previously shown that ERβ is expressed in intestinal epithelial at low levels, and has specific functions, whereas there is not sufficient support that ERα is expressed in the colon (Anderson et al, Nat Comm, 2017). While we did not sequence the colon transcriptome of DPN-treated mice, we evaluated the E2-mediated response of ERβ-target genes (as predicted from our ERβ chromatin immunoprecipitation (ChIP)-seq data from human CRC cell lines, with transduced ERβ, GSE149979) in the male colon transcriptome. This resulted in identification of 23 HFD and E2-regulated genes with cis-chromatin ERβ-binding sites (Fig. 4A). These included the cell cycle genes *Stk11* and *Wee1* and the inflammatory related gene *Anxa1* among others (Fig. 4B). Moreover, we compared the HFD regulated genes, which also presented sex-differences for potential E2 regulation in females. This comprised of 287 genes of which 45 showed ERβ binding sites, including the orphan nuclear receptor *Nr4a1 (Nur77*, Fig. 4A-B). qPCR analysis of colon tissue from the ligand treated mice corroborated the HFD-induced upregulation of *Anxa1* and that this was indeed attenuated by both E2 and DPN (ERβ) but not PPT in females, whereas it was not affected in males (Fig. 4H). Further, the HFD-induction of the M2 macrophage marker *Arg1* was also stronger in females, and this was blocked by both ERα and ERβ activation (Fig. 4H). In addition, we have previously shown that ERβ can cross talk and regulate NFκB signaling in both human colon cell lines and in the colon of ERβ intestinal knockout mice (Hases et al, under review). While the effect of deleted colonic ERβ on these genes was strongest in males, we here observe that HFD-induced expression of several NFκB target genes (*Cxcl5, Nos2)* were evident in females, and that this was blocked by both ERβ (DPN) and ERα (PPT) activation. We conclude that ERα systemically affects obesity, and that ERβ through local gene regulation downregulates proliferation and inflammatory genes (*Cxcl5* and *Nos2*) in the colon. This data provides evidence that estrogen through systemic and local effects via ERα and ERβ can attenuate inflammatory effects in the colon.

**Figure 4.**
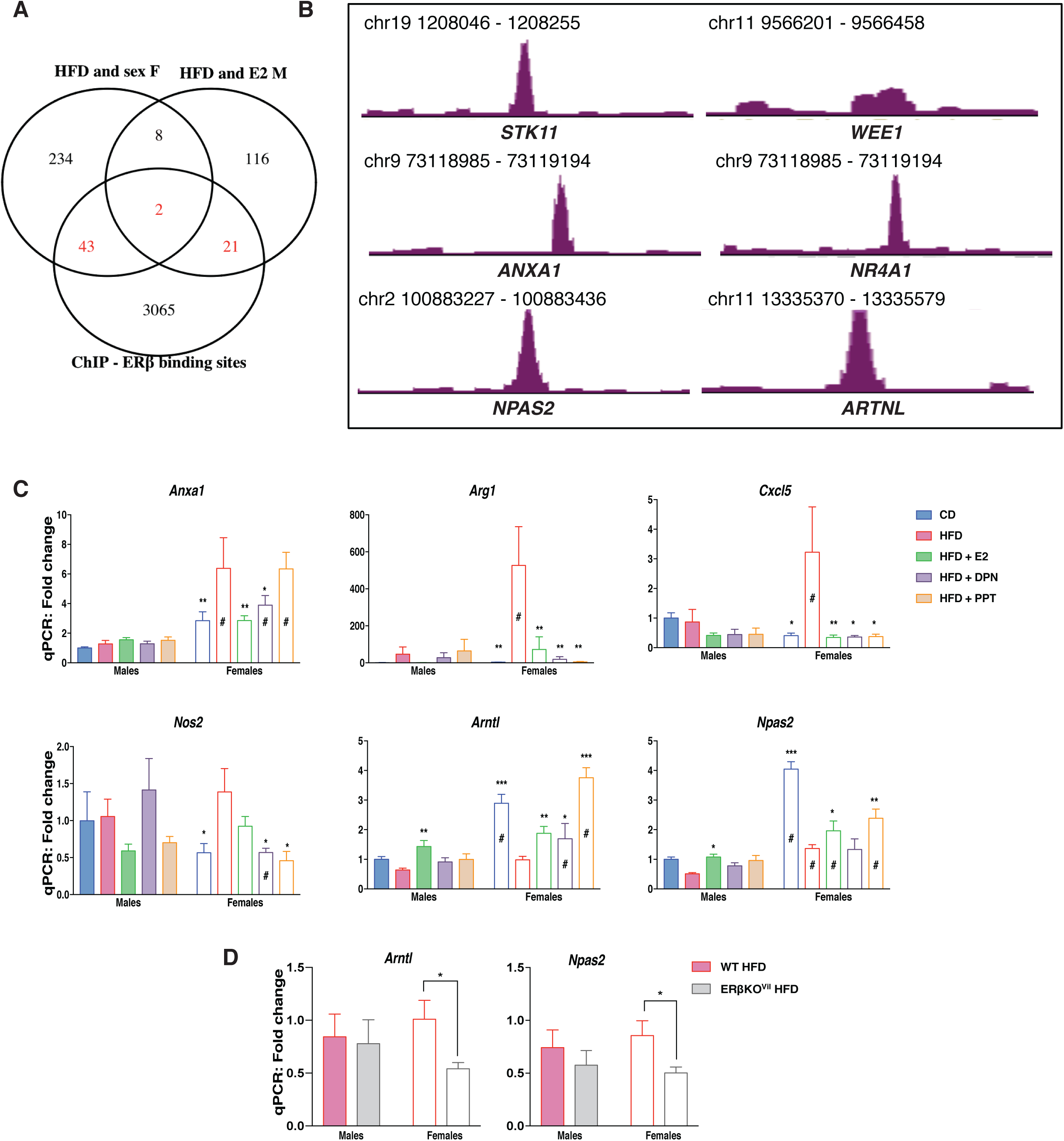
Estrogen through both ERα and ERβ can modulate the expression of specific sets of genes under HFD in both sexes. (A) Venn diagram of the E2 regulated genes under HFD in males and the HFD regulated genes in females compared to ERβ binding sites in two different CRC cell lines (SW480 and HT29). (B) ERβ binding sites revealed with ChIP-sequencing in CRC cell lines. (C) qPCR gene expression in the colon of female and male mice on a CD, HFD and HFD treated with E2, DPN and PPT (n=5-15). (D) qPCR gene expression in the colon of wild type and intestinal specific ERβ KO mice fed a HFD. The results are presented as mean ±SEM. Two-way ANOVA with uncorrected Fisher’s LSD test, *p<0.05, **p<0.01, ***p<0.001. # Indicate sex-differences.

### Sex, hormones, and diet influence regulation of circadian rhythm genes in colon

The circadian rhythm is essential for intestinal homeostasis. Disruptions can increase intestinal permeability, proliferation, exacerbate colitis, modulate microbiota composition, and impact the immune system (37-40). HFD is known to impact the circadian rhythmicity in both liver and adipose tissue, however very little is known about the impact of HFD of the circadian rhythm in the colon epithelium. Here we identified a sex difference in the expression of clock genes with higher expression of *Arntl, Npas2* and *Cry1*, in female colon and higher expression of *Per3* and *Nr1d1 (Rev-ErbA)* in male colon (Fig. 1F). *Arntl* and *Npas2* were downregulated and *Per3* upregulated by HFD in both sexes, whereas *Cry1* was downregulated by HFD specifically in females and *Per2* was upregulated by HFD specifically in males (Fig. 1F & 2A). In addition to exhibiting sex differences in their expression levels, ER ligands could oppose the HFD-induced downregulation of both *Arntl* and *Npas2*, in both sexes (Fig. 3D & 4C). ChIP-Seq indicated that ERβ is a direct regulator of the clock genes *Arntl* and *Npas2* (Fig. 4G). In males, both *Arntl* and *Npas2* were upregulated in the colon through estrogen, but significant regulation by ERβ (DPN) were not observed (Fig. 4H). However, we could confirm that the circadian clock gene *Arntl* indeed was upregulated by DPN (ERβ) and, also, by E2 and PPT (ERα) in females (Fig. 4H). These evidence points towards local regulation of clock genes by ERβ in the colon. The circadian rhythm of the colon has essential impacts on the body, and if this is impacted by ERβ this has substantial implications for our understanding of how hormones affect the body. In order to provide conclusive evidence of ERβ-mediated regulation of *Arntl* and *Npas2* in the *in vivo* colon, we investigated the impact of intestinal-specific deletion of ERβ in mice fed a HFD. Analyzing the colon gene expression in mice with and without intestinal ERβ after 13 weeks of HFD, using qPCR, we found a significant decrease in the expression of both *Arntl* and *Npas2* in the female knockout mice (Fig. 4D). Our results show, for the first time, that the expression of genes, including clock genes, involved in the circadian/rhythmic signaling pathway in colon is sex-dependent, regulated by HFD in both sexes upon HFD, and regulated by ERβ in the colon epithelial cells.

## Discussion

Our objective with this study was to determine whether estrogen could modulate the colon microenvironment during HFD and, if so, to dissect this mechanism and identify sex differences. Several studies have suggested that both ERα and ERβ can improve the metabolic syndrome, a risk factor for CRC (24-26). Other studies support a protective role of specifically ERβ against CRC (reviewed in (17)). The role of ERβ, or ERα, on the colon during HFD-induced obesity has, however, not been investigated. We here demonstrate that estrogen indeed modulates the colon microenvironment in mice fed HFD, and that ERα- and ERβ-selective ligands can counteract different facets of HFD-induced phenotypes, including at the level of colonic gene expression.

Looking at the metabolic data, males appeared more sensitive to HFD, while E2 treatment as well as ERα-specific activation by PPT improved the glucose metabolisms in males, in line with previous studies (34, 35). Some studies have also reported a protective effect of ERβ against HFD-induced obesity, but the data are still conflicting. Foryst-Ludwig and colleagues found that ERβ protected against HFD-induced weight gain, but still elicited pro-diabetogenic effects by impairing insulin sensitivity and glucose tolerance (41). We could not confirm that ERβ protected against HFD-induced weight gain however, in accordance with Foryst-Ludwig *et al*., we found that its activation impaired glucose levels in females (Suppl. fig. 1C) and not in males (Fig. 3A).

Our findings detail several other significant sex differences. In particular, there was a notable sex difference in the colon transcriptome in mice under control diet setting, which was not expected. However, the diets did not contain soy or any phytoestrogens compared to a standardized chow diet, thus we could observe the impact of endogenous estrogen signaling without external interference apart from the ligand treatments. This condition may have contributed to increased sex differences compared to a regular chow, which contains high levels of soy (phytoestrogen), which would activate estrogen signaling in both males and females. The sex-differences observed in both dietary conditions (CD and HFD) were related to hypoxia, immune response, cell proliferation, circadian rhythm and Wnt signaling. Thus, our data bring in a role of sex hormones in the basal regulation of these pathways in colon. Intriguingly, we found that estrogen treatment in males indeed regulated the immune response, cell proliferation and rhythmic processes in colonic tissue.

Interestingly, HFD-induced pro-inflammatory modifications occur earlier in the colon than in the adipose tissue. These changes are characterized by increased pro-inflammatory macrophages infiltration and epithelia permeability in the colon, which eventually can lead to insulin resistance (7). Estrogen treatment has been shown to elicit an anti-inflammatory response in the colon of mouse models for colitis (28, 46). Our transcriptomic data revealed that while both sexes showed altered immune response upon HFD feeding, there were sex differences in the immune response. This was supported at the histological level by a significant increase of F4/80^+^ macrophage infiltration in the colon epithelium upon HFD in both sexes, but with a significantly higher overall count of F4/80^+^ macrophages in females. The F4/80^+^ macrophage infiltration was significantly reduced by ERβ activation (DPN). In addition, our mechanistic data from ChIP-Seq in CRC cell lines demonstrated that ERβ could bind to cis-regulatory chromatin regions of both *Anxa1* and the orphan nuclear receptor *Nr4a1 (Nur77). Anxa1* is known to recruit monocytes (42) and *Nur77* can regulate the immune system and can suppress colitis induced with DSS (43). Additionally, in females, we found that ERβ activation could modulate the expression of *Arg1* and *Nos2*, markers for M2 anti-inflammatory and M1 pro-inflammatory macrophages, respectively (44, 45). Thus, our study delineates how ERβ contributes to anti-inflammatory effects of estrogen in the *in vivo* colon.

Furthermore, we have recently shown in colitis-induced CRC mouse model and colonic human cells that ERβ expressed in intestinal epithelial cells can down regulate specifically TNFα/NFkB activation network (Hases et al, under review). ERβ is not detected in macrophages, while it is expressed in intestinal epithelial cells (see Human Protein Atlas). However, crosstalk between the colonic epithelial cells and macrophages has been demonstrated in IBD (47) and with HFD in an obesity context (7). Although the etiology of obesity and IBD are different, both are characterized by gut inflammation with common inflammatory pathways including TNFα, IL6 and IL1b (48-51). Thus, our data suggest that specific activation of intestinal ERβ may have a beneficial impact to modulate gut inflammation in both pathologies.

Additionally, the colon transcriptome responded in a sex-dependent manner upon HFD-feeding. The transcriptomic analysis revealed male-enriched pathways involved in cell proliferation, e.g. cell cycle, DNA replication and proliferation. In addition, E2 treatment in males opposed the HFD-induced effect on cell cycle and proliferation. This was corroborated at the functional level with increased Ki67 cell proliferation upon HFD specifically in males. Possibly, the females in our study were already protected against HFD-induced cell proliferation by their endogenous estrogen. Obesity is a well-known risk factor for CRC, and HFD-induced obesity exacerbates colon tumor development in mice treated with azoxymethane by enhancing the colonic cell proliferation (52). Previous *in vitro* results have demonstrated anti-proliferative effects by exogenously expressed ERβ in CRC cell lines (53-55). In line with these data, we found a significant decrease in HFD-induced cell proliferation upon ERβ activation (DPN). These results demonstrate that ERβ expression in the colon epithelial cells can protect against the HFD-induced cell proliferation, and potentially undermine the promotion of colon tumor development.

The most unexpected finding was that key circadian clock genes (*Npas2, Arntl (Bmal1), Per3)* were differentially expressed between the sexes in the colon, during both dietary conditions. The circadian clock regulates critical processes involved in inflammation and proliferation. HFD feeding is known to impact the circadian rhythm in the hypothalamus, liver and fat tissue, and to disrupt the eating behavior. However timed HFD-feeding can reset the circadian metabolism and help prevent obesity and metabolic disorders (56, 57), indicating a major contribution of the circadian rhythm in the pathogenesis of the disease. The circadian rhythm has been shown to be dysregulated in both obesity and IBD (58, 59). The circadian clock is regulated by different factors including sex hormones such as estrogen (E2) and there are evidences of sex-dependent regulation of the circadian clock (60). Palmisano *et al*. have shown that females are protected against HFD-induced circadian disruption in the liver when fed a 45% HFD for 8 weeks, indicating a role for estrogen in regulation of the circadian rhythm (62). Further, in breast cancer cell lines, ERα was shown to increase the protein and mRNA expression of Clock (61), indicating the potential of estrogen receptors to regulate clock genes at the tissue level. However the impact of HFD and estrogen on the circadian rhythm in the colon epithelium has not previously been investigated. Here we show for the first time that in mice fed HFD, both sexes present an altered expression of *Npas2* and *Arntl (Bmal1)* in colon. We also found that ligand activation of estrogen signaling can modulate the expression of *Npas2* and *Arntl (Bmal1)* in the colon. Interestingly, our mechanistic data from ChIP-Seq in CRC cell lines demonstrated that ERβ could bind to cis-regulatory chromatin regions of both *Npas2* and *Arntl (Bmal1)*. This suggests that there are local effects acting via ERβ in the colon, and we could confirm this using our intestinal-specific ERβ knockout mice. The expression of *Npas2*, which was enhanced by DPN treatment (specific-ERβ activation) in males, was indeed significantly decreased by ERβ knockout in the colon epithelium of female mice on a HFD.

Worth noting, is that ERα activation by PPT also regulated the expression of *Npas2* and *Arntl* in the colon. However, available data do not support significant expression of ERα in the colon (63), and we therefore assume that PPT effects are more global through its effect on the overall body’s metabolism. However, we cannot completely exclude local effects of ERα in colon, in particular via the immune cells. We did notice a two-fold increase of colonic ERα mRNA levels in females upon HFD-feeding. Both receptors bind estrogen response element (ERE) in the chromatin, but can otherwise differ in their protein interactions. The ERβ-binding site by the *Arntl* gene included an ERE, which ERα is also likely to bind, if expressed. However, the *Npas2* ERβ-binding site was indicated to be through tethering with JUN/AP-1, and is likely to be specific for ERβ.

It should be emphasized that although all mice were consistently sacrificed at the same time of the day, this was not a circadian experiment and we did not investigate different zeitgeber time points. Hence, while our results highlight the implication of ERβ in the regulation of core clock genes in the colon, further experiments are needed to conclude its impact on their rhythmicity and whether this is different in male and female colon.

In conclusion, our data provide evidence that the colon microenvironment responds to HFD in both sexes, but in a sex-dependent manner, as illustrated in Figure 5. Both sexes present an altered expression of core clock genes and increased macrophage infiltration, whereas males showed a significant increase of epithelial cell proliferation. All of the above have been shown to play important roles in the pathogenesis of HFD-induced metabolic disorders and colitis. Entirely novel findings in our study demonstrate that estrogen signaling can modulate the colon microenvironment during HFD. In particular, the expression of the core clock gene *Npas2*, as well as the colonic cell proliferation and macrophage infiltration can be modulated by estrogen via ERβ in a sex-dependent manner. A part of this effect may be triggered by local activation of ERβ in the colon, as suggested by the ChIP-seq data. And we demonstrate using our intestinal-specific ERβ knockout mice that ERβ expression in the colon is a direct regulator of both *Arntl* and *Npas2*. This study opens up new insights, where ERβ exhibits beneficial action against deleterious effects of diet-induced obesity on colon microenvironment.

**Figure 5.**
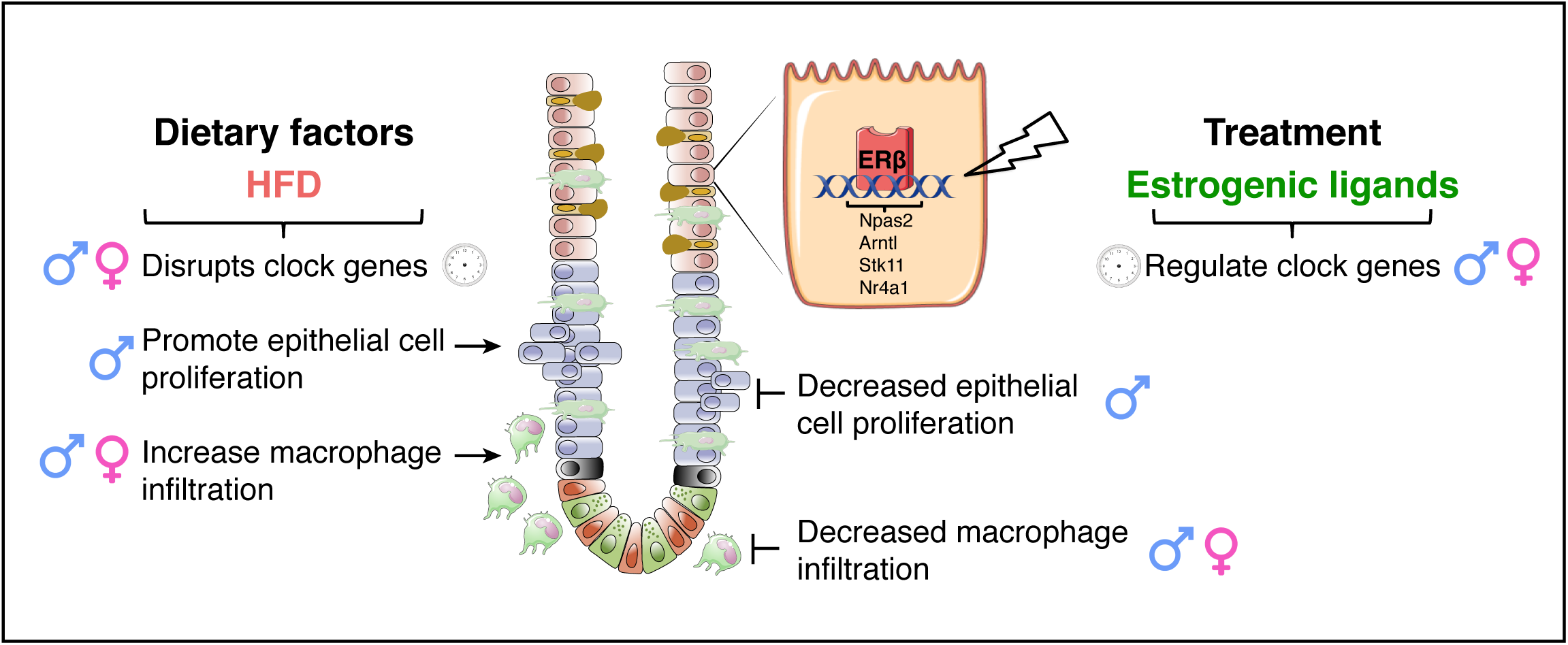
Schematic illustration; proposed model for estrogen regulation of the colon microenvironment during HFD-induced obesity. The HFD-induced impaired colonic circadian rhythm and increased cell proliferation was opposed by ERβ activation with DPN in males. The HFD-induced increase in macrophage infiltration was repressed by ERβ activation in both females and males.

## Material and Methods

### Animals

Five to six-week old male and female C57BL/6J mice were obtained from in-house breeding. Mice lacking ERβ specifically in the epithelial cells in the intestine were generated by crossing ERβ^flox/flox^ mice (B6.129×1-Esr2^tm1Gust^, Jan-Åke Gustafsson’s laboratory) with the transgenic mice bearing cre-recombinase expressed under the control of the enterocyte-specific Villin promoter (B6.SJL-Tg(Vil-cre)997 Gum/J; Jackson Laboratory, Bar Harbor, ME). Littermates (ERβ^flox/flox^) lacking the Cre allele were used as controls (referred to as WT). The genotype was confirmed with standard PCR protocol. Animals were housed under controlled environment at 20°C with a 12-h light-dark cycle. The animals were fed HFD (D12492, 60% kcal fat, Research Diet) or a control diet corresponding to a matched low-fat diet (D12450J, 10% kcal fat, Research Diet) and water was provided *ad libitum*. For the treatment with estrogenic ligands, at 10 weeks of diet the mice were injected *i*.*p*. every other day for a total of 9 injections with 0.05 or 0.5mg/kg BW for 17β-estradiol (E2, Sigma-Aldrich), 2.5 mg/kg BW for 4,4’,4’’-(4-Propyl-[1H]-pyrazole-1,3,5-triyl)trisphenol (PPT, Tocris), 5 mg/kg BW for 2,3-*bis*(4-Hydroxyphenyl)-propionitrile (DPN, Tocris), or vehicle. The ligands were prepared in a solution of 40% PEG400, 5% DMSO and 55% water. The local ethical committee of the Swedish National Board of Animal Research (N230/15) approved all experimental protocols.

## Material and Methods

### Animals

Five to six-week old male and female C57BL/6J mice were obtained from in-house breeding. Mice lacking ERβ specifically in the epithelial cells in the intestine were generated by crossing ERβ^flox/flox^ mice (B6.129×1-Esr2^tm1Gust^, Jan-Åke Gustafsson’s laboratory) with the transgenic mice bearing cre-recombinase expressed under the control of the enterocyte-specific Villin promoter (B6.SJL-Tg(Vil-cre)997 Gum/J; Jackson Laboratory, Bar Harbor, ME). Littermates (ERβ^flox/flox^) lacking the Cre allele were used as controls (referred to as WT). The genotype was confirmed with standard PCR protocol. Animals were housed under controlled environment at 20°C with a 12-h light-dark cycle. The animals were fed HFD (D12492, 60% kcal fat, Research Diet) or a control diet corresponding to a matched low-fat diet (D12450J, 10% kcal fat, Research Diet) and water was provided *ad libitum*. For the treatment with estrogenic ligands, at 10 weeks of diet the mice were injected *i*.*p*. every other day for a total of 9 injections with 0.05 or 0.5mg/kg BW for 17β-estradiol (E2, Sigma-Aldrich), 2.5 mg/kg BW for 4,4’,4’’-(4-Propyl-[1H]-pyrazole-1,3,5-triyl)trisphenol (PPT, Tocris), 5 mg/kg BW for 2,3-*bis*(4-Hydroxyphenyl)-propionitrile (DPN, Tocris), or vehicle. The ligands were prepared in a solution of 40% PEG400, 5% DMSO and 55% water. The local ethical committee of the Swedish National Board of Animal Research (N230/15) approved all experimental protocols.

### Fasting glucose

Mice were fasted for 6h prior the measurement of the glucose level. Briefly, a small piece of the tail was cut with a scalpel, a drop of blood was removed, and glucose level immediately measured with a OneTouch Ultra glucometer (AccuChek Sensor, Roche Diagnostics).

### Tissue collection

Colons were harvested, washed, measured and opened along the long axis. The colons were then either embedded in OCT (Tissue-tek, Sakura) and prepared for immunohistochemistry (IHC) analyses using the “swiss roll” method, or snap frozen in liquid nitrogen for further analysis.

### Fat quantification

Mice were dissected and fat pads from the abdominal and posterior subcutaneous regions carefully removed and weighted. Total fat (TF) was calculated as the sum of all fat depots including gonadal fat, retroperitoneal fat, omental fat and inguinal/gluteal fat depots. Visceral adipose (VAT) comprised of the sum of gonadal fat, retroperitoneal fat and omental fat. The subcutaneous fat (SAT) comprised of the inguinal/gluteal fat depots.

### ELISA

Estradiol levels were measured in mouse serum by mouse/rat estradiol kit (ES180S-100, Calbiotech) according to manufacturer’s instructions. Serum samples were diluted 1:5 for vehicle treated CD and HFD and 1:12 for E2 treated mice with assay diluent. Calculations were performed using standard controls provided with the kit.

### Immunohistochemistry

Colonic samples embedded in OCT were cryo-sectioned at 8µm thickness. Sections were fixed in 4% formaldehyde for 10 min and endogenous peroxidase activity was blocked with 3% hydrogen peroxidase in 50 % methanol and PBS for 30 min. Slides were blocked with 5% normal goat serum for 30 min at 4°C and blocked for unspecific avidin/biotin binding (DAKO). The sections were incubated overnight at 4°C with the primary antibodies, anti-F4/80 (1:100) (MCA497R, Biorad), anti-Ki67 (1:100) (SP6, Invitrogen) in 0.1 % IGEPAL in PBS. Negative controls without primary antibodies were used for each slide. The sections were incubated with appropriate biotin conjugated anti IgG secondary antibodies (1:500) for 1h at room temperature (RT), and then incubated with avidin-biotin complex (ABC) (Thermofisher) for 1h at RT. Sections were developed with Liquid DAB+ (3,3-diaminobenzidine) Substrate Chromogen System (DAKO) and counterstained with Mayer’s Hematoxylin (Sigma-Aldrich). After dehydration in ethanol and xylene, the slides were cover-slipped using Pertex (Histolab). Ten to fifteen random images from three fully stained sections per animal were randomly taken using a BX53 light microscope and CAM-SC50 camera (Olympus). Positive stained cells per crypts were counted using QuPath and approximately 30 crypts per animal were analyzed.

### RNA isolation and quantitative PCR

Frozen colon distal tissue and colon epithelial layer were homogenized with a tissue lyser (Qiagen, Chatsworth, CA). Total RNA was isolated with QIAzol and purified using miRNeasy Mini Kit (Qiagen, Chatsworth, CA) according to the standard protocol and on-column DNAse treatment was applied. Quantitative and qualitative analyses of the RNA were performed with NanoDrop 1000 spectrophotometer and Agilent Tapestation 2200 (Agilent Technologies, Palo Alto, CA), respectively. All samples had RNA integrity >6.5. One microgram RNA was reversed transcribed using iScript cDNA synthesis kit (Biorad) according to standard protocol. Ten nanogram of cDNA were used to perform qPCR in the CFX96 Touch System (Biorad), with iTaq universal SYBR Green supermix (Biorad) as recommended by the supplier. Samples were run in duplicates and the relative gene expression was calculated as the mean per group using the ΔΔCt method, normalized to the geometric mean of three reference genes (*Actb, Eef2* and *Tbp*). Primer sequences are provided in supplementary table 1.

### RNA-seq

RNA from colon tissue was prepared from 6 biological replicates, from male and female fed with CD and HFD, and males fed with HFD and treated with E2 (0.5mg/kg BW). Sequencing library preparation with Illumina RiboZero and sequencing using Illumina NovaSeq6000 was performed at Sweden’s National Genomics Infrastructure (NGI). At least 17M paired-end reads (2×51bp in length) were generated for each sample, and mapped against mouse genome (GRCm38) using STAR. FeatureCounts and StringTie were used to generate gene counts and TPM values. DESeq2 was used to calculate differentially expressed genes (DEG) with raw counts as input and the Benjamini-Hochberg procedure was used to estimate FDR. Genes were considered as significantly differentially expressed if P-value<0.05 and log2FC>|0.4|. Gene ontology/biological function was performed with DAVID bioinformatics website.

### Statistical analysis

GraphPad Prism was used for statistical analysis (GraphPad Software Inc, La Jolla, CA). The results are presented as mean ±SEM. A (two-tailed) Welch’s t-test was used for comparison between two groups. One-way analysis of variance (ANOVA) was used for comparison between multiple conditions followed by fisher’s LSD test. Two-way ANOVA was used for comparison between multiple conditions in two different groups followed by fisher’s LSD test. A p-value <0.05 was considered being statistically significant (* p<0.05, ** p<0.01, *** p<0.001).

## Acknowledgements

The authors acknowledge support from the National Genomics Infrastructure in Stockholm funded by Science for Life Laboratory, the Knut and Alice Wallenberg Foundation and the Swedish Research Council, and SNIC/Uppsala Multidisciplinary Center for Advanced Computational Science for assistance with massively parallel sequencing and access to the UPPMAX computational infrastructure. Additionally, Viktor Jonsson from NBIS/SciLifeLab advised on the RNA-seq analysis.

## Author contributions

L.H. and A.A. designed the *in vivo* study; L.H., A.A., R.I., M.B., C.S. and M.K-A performed all the *in vivo* experiments; A.A. and M.B. performed the histology experiments; R.I. performed ChIP experiments; L.H., and A.A. performed the qPCR experiments; L.H. performed the RNA-seq analyses; L.H., A.A. and C.W. interpreted results; L.H., R.I., and A.A. prepared figures; L.H., A.A. and C.W wrote manuscript; M.K-A. advised on the metabolic parameters; A.A. and C.W. initiated, coordinated the study and supervised. All authors discussed the results and approved final version of manuscript.

## Data and materials availability

Gene expression data are deposited in the NCBI Gene Expression Omnibus database [GSE149811].

## References

1. D. P. Guh et al., The incidence of co-morbidities related to obesity and overweight: a systematic review and meta-analysis. BMC Public Health 9, 88 (2009).

2. K. C. Harris, L. K. Kuramoto, M. Schulzer, J. E. Retallack, Effect of school-based physical activity interventions on body mass index in children: a meta-analysis. Cmaj 180, 719–726 (2009).

3. Z. Dai, Y. C. Xu, L. Niu, Obesity and colorectal cancer risk: a meta-analysis of cohort studies. World J Gastroenterol 13, 4199–4206 (2007).

4. A. A. Moghaddam, M. Woodward, R. Huxley, Obesity and risk of colorectal cancer: a meta-analysis of 31 studies with 70,000 events. Cancer Epidemiol Biomarkers Prev 16, 2533–2547 (2007).

5. L. Pietrzyk, A. Torres, R. Maciejewski, K. Torres, Obesity and Obese-related Chronic Low-grade Inflammation in Promotion of Colorectal Cancer Development. Asian Pac J Cancer Prev 16, 4161–4168 (2015).

6. M. Bardou, A. N. Barkun, M. Martel, Obesity and colorectal cancer. Gut 62, 933–947 (2013).

7. Y. Kawano et al., Colonic Pro-inflammatory Macrophages Cause Insulin Resistance in an Intestinal Ccl2/Ccr2-Dependent Manner. Cell Metab 24, 295–310 (2016).

8. S. Beyaz et al., High-fat diet enhances stemness and tumorigenicity of intestinal progenitors. Nature 531, 53–58 (2016).

9. H. Luck et al., Regulation of obesity-related insulin resistance with gut anti-inflammatory agents. Cell Metab 21, 527–542 (2015).

10. P. D. Cani et al., Changes in gut microbiota control metabolic endotoxemia-induced inflammation in high-fat diet-induced obesity and diabetes in mice. Diabetes 57, 1470–1481 (2008).

11. M. A. Hildebrandt et al., High-fat diet determines the composition of the murine gut microbiome independently of obesity. Gastroenterology 137, 1716-1724.e1711-1712 (2009).

12. M. Serino et al., Metabolic adaptation to a high-fat diet is associated with a change in the gut microbiota. Gut 61, 543–553 (2012).

13. K. A. Kim, W. Gu, I. A. Lee, E. H. Joh, D. H. Kim, High fat diet-induced gut microbiota exacerbates inflammation and obesity in mice via the TLR4 signaling pathway. PLoS One 7, e47713 (2012).

14. Z. Liu et al., Diet-induced obesity elevates colonic TNF-alpha in mice and is accompanied by an activation of Wnt signaling: a mechanism for obesity-associated colorectal cancer. J Nutr Biochem 23, 1207–1213 (2012).

15. H. Kim, E. L. Giovannucci, Sex differences in the association of obesity and colorectal cancer risk. Cancer Causes Control 28, 1–4 (2017).

16. D. Zheng et al., Sexual dimorphism in the incidence of human cancers. BMC Cancer 19, 684 (2019).

17. C. Williams, A. DiLeo, Y. Niv, J. A. Gustafsson, Estrogen receptor beta as target for colorectal cancer prevention. Cancer Lett 372, 48–56 (2016).

18. R. A. Lobo, Hormone-replacement therapy: current thinking. Nat Rev Endocrinol 13, 220–231 (2017).

19. E. Botteri et al., Menopausal hormone therapy and colorectal cancer: a linkage between nationwide registries in Norway. BMJ Open 7, e017639 (2017).

20. J. Simin, R. Tamimi, J. Lagergren, H. O. Adami, N. Brusselaers, Menopausal hormone therapy and cancer risk: An overestimated risk? Eur J Cancer 84, 60–68 (2017).

21. G. Stachowiak, T. Pertynski, M. Pertynska-Marczewska, Metabolic disorders in menopause. Prz Menopauzalny 14, 59–64 (2015).

22. S. R. Salpeter et al., Meta-analysis: effect of hormone-replacement therapy on components of the metabolic syndrome in postmenopausal women. Diabetes Obes Metab 8, 538–554 (2006).

23. A. Bouman, M. J. Heineman, M. M. Faas, Sex hormones and the immune response in humans. Hum Reprod Update 11, 411–423 (2005).

24. P. A. Heine, J. A. Taylor, G. A. Iwamoto, D. B. Lubahn, P. S. Cooke, Increased adipose tissue in male and female estrogen receptor-alpha knockout mice. Proc Natl Acad Sci U S A 97, 12729–12734 (2000).

25. M. Yepuru et al., Estrogen receptor-{beta}-selective ligands alleviate high-fat diet- and ovariectomy-induced obesity in mice. J Biol Chem 285, 31292–31303 (2010).

26. M. Gonzalez-Granillo et al., Selective estrogen receptor (ER)beta activation provokes a redistribution of fat mass and modifies hepatic triglyceride composition in obese male mice. Mol Cell Endocrinol 502, 110672 (2019).

27. V. Giroux, G. Bernatchez, J. C. Carrier, Chemopreventive effect of ERbeta-Selective agonist on intestinal tumorigenesis in Apc(Min/+) mice. Molecular carcinogenesis 50, 359–369 (2011).

28. H. J. Son et al., Effect of Estradiol in an Azoxymethane/Dextran Sulfate Sodium-Treated Mouse Model of Colorectal Cancer: Implication for Sex Difference in Colorectal Cancer Development. Cancer Res Treat 51, 632–648 (2019).

29. C. H. Song et al., Effects of 17beta-Estradiol on Colonic Permeability and Inflammation in an Azoxymethane/Dextran Sulfate Sodium-Induced Colitis Mouse Model. Gut Liver 12, 682–693 (2018).

30. C. H. Song et al., Effects of 17beta-estradiol on colorectal cancer development after azoxymethane/dextran sulfate sodium treatment of ovariectomized mice. Biochem Pharmacol 164, 139–151 (2019).

31. D. Saleiro et al., Estrogen receptor-beta protects against colitis-associated neoplasia in mice. International journal of cancer 131, 2553–2561 (2012).

32. V. Giroux, F. Lemay, G. Bernatchez, Y. Robitaille, J. C. Carrier, Estrogen receptor beta deficiency enhances small intestinal tumorigenesis in ApcMin/+ mice. Int J Cancer 123, 303–311 (2008).

33. M. Gonzalez-Granillo et al., Sex-specific lipid molecular signatures in obesity-associated metabolic dysfunctions revealed by lipidomic characterization in ob/ob mouse. Biol Sex Differ 10, 11 (2019).

34. U. S. Pettersson, T. B. Walden, P. O. Carlsson, L. Jansson, M. Phillipson, Female mice are protected against high-fat diet induced metabolic syndrome and increase the regulatory T cell population in adipose tissue. PLoS One 7, e46057 (2012).

35. R. S. Dakin, B. R. Walker, J. R. Seckl, P. W. Hadoke, A. J. Drake, Estrogens protect male mice from obesity complications and influence glucocorticoid metabolism. Int J Obes (Lond) 39, 1539–1547 (2015).

36. E. Riant et al., Estrogens protect against high-fat diet-induced insulin resistance and glucose intolerance in mice. Endocrinology 150, 2109–2117 (2009).

37. R. Pagel et al., Circadian rhythm disruption impairs tissue homeostasis and exacerbates chronic inflammation in the intestine. Faseb j 31, 4707–4719 (2017).

38. O. O. Kyoko et al., Expressions of tight junction proteins Occludin and Claudin-1 are under the circadian control in the mouse large intestine: implications in intestinal permeability and susceptibility to colitis. PLoS One 9, e98016 (2014).

39. J. A. Deaver, S. Y. Eum, M. Toborek, Circadian Disruption Changes Gut Microbiome Taxa and Functional Gene Composition. Front Microbiol 9, 737 (2018).

40. R. M. Voigt et al., Circadian disorganization alters intestinal microbiota. PLoS One 9, e97500 (2014).

41. A. Foryst-Ludwig et al., Metabolic actions of estrogen receptor beta (ERbeta) are mediated by a negative cross-talk with PPARgamma. PLoS Genet 4, e1000108 (2008).

42. S. McArthur et al., Definition of a Novel Pathway Centered on Lysophosphatidic Acid To Recruit Monocytes during the Resolution Phase of Tissue Inflammation. J Immunol 195, 1139–1151 (2015).

43. A. A. Hamers et al., Deficiency of Nuclear Receptor Nur77 Aggravates Mouse Experimental Colitis by Increased NFkappaB Activity in Macrophages. PLoS One 10, e0133598 (2015).

44. P. Das, A. Lahiri, A. Lahiri, D. Chakravortty, Modulation of the arginase pathway in the context of microbial pathogenesis: a metabolic enzyme moonlighting as an immune modulator. PLoS Pathog 6, e1000899 (2010).

45. M. Munder, K. Eichmann, M. Modolell, Alternative metabolic states in murine macrophages reflected by the nitric oxide synthase/arginase balance: competitive regulation by CD4+ T cells correlates with Th1/Th2 phenotype. J Immunol 160, 5347–5354 (1998).

46. C. H. Song et al., Effects of 17β-estradiol on colorectal cancer development after azoxymethane/dextran sulfate sodium treatment of ovariectomized mice. Biochem Pharmacol 164, 139–151 (2019).

47. S. Al-Ghadban, S. Kaissi, F. R. Homaidan, H. Y. Naim, M. E. El-Sabban, Cross-talk between intestinal epithelial cells and immune cells in inflammatory bowel disease. Sci Rep 6, 29783 (2016).

48. S. Ding et al., High-fat diet: bacteria interactions promote intestinal inflammation which precedes and correlates with obesity and insulin resistance in mouse. PLoS One 5, e12191 (2010).

49. M. Gulhane et al., High Fat Diets Induce Colonic Epithelial Cell Stress and Inflammation that is Reversed by IL-22. Sci Rep 6, 28990 (2016).

50. P. A. Kern, S. Ranganathan, C. Li, L. Wood, G. Ranganathan, Adipose tissue tumor necrosis factor and interleukin-6 expression in human obesity and insulin resistance. Am J Physiol Endocrinol Metab 280, E745–751 (2001).

51. H. C. Reinecker et al., Enhanced secretion of tumour necrosis factor-alpha, IL-6, and IL-1 beta by isolated lamina propria mononuclear cells from patients with ulcerative colitis and Crohn’s disease. Clin Exp Immunol 94, 174–181 (1993).

52. I. Tuominen et al., Diet-induced obesity promotes colon tumor development in azoxymethane-treated mice. PLoS One 8, e60939 (2013).

53. K. Edvardsson, A. Strom, P. Jonsson, J. A. Gustafsson, C. Williams, Estrogen receptor beta induces antiinflammatory and antitumorigenic networks in colon cancer cells. Mol Endocrinol 25, 969–979 (2011).

54. K. Edvardsson et al., Estrogen receptor beta expression induces changes in the microRNA pool in human colon cancer cells. Carcinogenesis 34, 1431–1441 (2013).

55. T. Nguyen-Vu et al., Estrogen receptor beta reduces colon cancer metastasis through a novel miR-205 - PROX1 mechanism. Oncotarget 7, 42159–42171 (2016).

56. A. Kohsaka et al., High-fat diet disrupts behavioral and molecular circadian rhythms in mice. Cell Metab 6, 414–421 (2007).

57. H. Sherman et al., Timed high-fat diet resets circadian metabolism and prevents obesity. Faseb j 26, 3493–3502 (2012).

58. A. Sobolewska-Włodarczyk et al., Circadian rhythm abnormalities - Association with the course of inflammatory bowel disease. Pharmacol Rep 68, 847–851 (2016).

59. O. Froy, Circadian rhythms and obesity in mammals. ISRN Obes 2012, 437198 (2012).

60. L. Yan, R. Silver, Neuroendocrine underpinnings of sex differences in circadian timing systems. J Steroid Biochem Mol Biol 160, 118–126 (2016).

61. L. Xiao et al., Induction of the CLOCK gene by E2-ERα signaling promotes the proliferation of breast cancer cells. PLoS One 9, e95878 (2014).

62. B. T. Palmisano, J. M. Stafford, J. S. Pendergast, High-Fat Feeding Does Not Disrupt Daily Rhythms in Female Mice because of Protection by Ovarian Hormones. Front Endocrinol (Lausanne) 8, 44 (2017).

63. M. Uhlén et al., Proteomics. Tissue-based map of the human proteome. Science 347, 1260419 (2015).

