## Supplementary information for "High-fat diet impacts the colon and its transcriptome in a sex-dependent manner that is modifiable by estrogens"

Linnea Hases[1]¶, Amena Archer¶, Rajitha Indukuri , Madeleine Birgersson, Christina Savva, Marion Korach-André, Cecilia Williams

**Corresponding author:** Cecilia Williams

### This PDF file includes:

Figure S1. Sex-differences in fat distribution and effects of estrogenic treatments in females.
Table S1. Primer sequences.

### Supplementary Figures

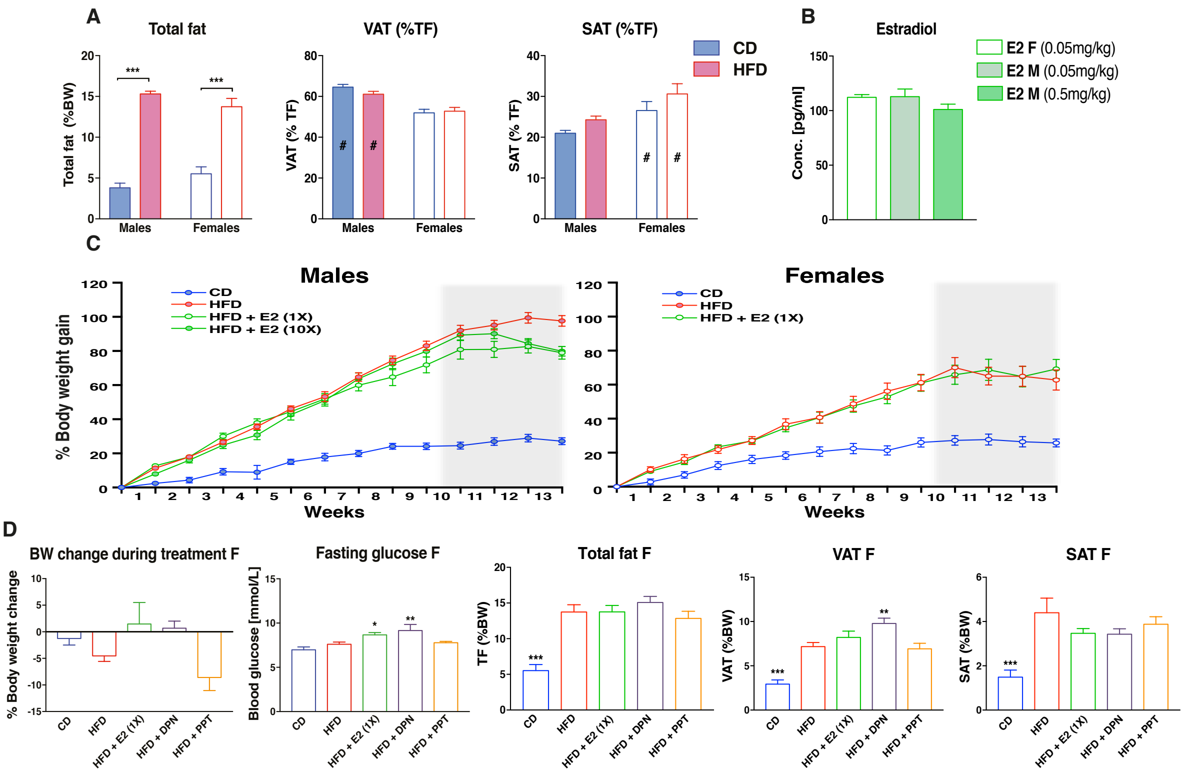

**Figure S1. Sex-differences in fat distribution and effects of estrogenic treatments in females. (A)** Total fat, VAT and SAT reported to total fat for males and females fed a CD or HFD. **(B)** Serum estradiol levels for females and males treated with estrogen (E2). **(C)** BW curves for males and females fed CD or HFD for a 13-weeks period and treated with E2 or vehicle for the last 3 weeks (highlighted in grey). **(D)** BW change during treatment, fasting glucose levels, total fat, VAT and SAT for females under a CD or HFD and treated with vehicle or different estrogenic ligands. The results are presented as mean ±SEM, (n=5-15). One-way and two-way ANOVA with uncorrected Fisher's LSD test, *p<0.05, **p<0.01, ***p<0.001. # Indicate sex-differences.

### Supplementary Tables

#### Table S1. Primer sequences.

| Gene | Forward primer (5' - 3') | Reverse primer (5' - 3') |
|---|---|---|
| **Genotyping primers** | | |
| Villin Cre | CAA GCC TGG CTC GAC GGC C | |
| Cre -50 antisense | ATC GAC CGG TAA TGC AGG CA | |
| ISP (loxP site) | TAG GGT ATG TTA TGT CAT GA | |
| BIASP (loxP site) | GTG GAT GCC TAT GAT CAC TGT GGA | |
| **qPCR primers** | | |
| mAnxa1 | AAGGTGGTCCTGGGTCAGC | TGAGCATTGGTCCTCTTGGT |
| mArg1 | CTCCAAGCCAAAGTCCTTAGAG | GGAGCTGTCATTAGGGACATCA |
| mArntl(Bmal1) | ACATAGGACACCTCGCAGAA | AACCATCGACTTCGTAGCGT |
| mB-actin | GGCTGTATTCCCCTCCATCG | CCAGTTGGTAACAATGCCATGT |
| mCxcl5 | GTTCCATCTCGCCATTCATGC | GCGGCTATGACTGAGGAAGG |
| mEef2 | CATCCTTGCGAGTGTCAGTGA | TGTCAGTCATCGCCCATGTG |
| mNos2 | CAGCTGGGVTGTACAAACCTT | CATTGGAAGTGAAGCGTTTCG |
| mNpas2 | ACGCAGATGTTCGAGTGGAAA | CGCCCATGTCAAGTGCATT |
| mTbp | GCAGCAAATCGCTTGGGATTA | ACCGTGAATCTTGGCTGTAAAC |
